# Computed Tomography-Derived Elastic Modulus as a Noninvasive Marker of Aortic Wall Integrity: Correlation with Histopathology in the Ascending Aorta

**DOI:** 10.1101/2025.06.27.662075

**Authors:** Fumio Yamana, Kazuo Shimamura, Takuji Kawamura, Takashi Shirakawa, Junki Yokota, Kansuke Kido, Ryoto Sakaniwa, Shunsuke Saito, Kizuku Yamashita, Akima Harada, Yoshiki Watanabe, Satoshi Sakakibara, Daisuke Yoshioka, Eiichi Morii, Shigeru Miyagawa

**Author notes:** Corresponding author: Kazuo Shimamura, Department of Cardiovascular Surgery, Osaka University Graduate School of Medicine, Osaka, Japan, 2-2 Yamadaoka, Suita, Osaka 565-0871, Japan.; E–mail Shigeru Miyagawa, Department of Cardiovascular Surgery, Osaka University Graduate School of Medicine, Osaka, Japan, 2-2 Yamadaoka, Suita, Osaka 565-0871, Japan.; E–mail.

## Abstract

**Background:** Ascending aortic aneurysms and dissections are life-threatening conditions often requiring prophylactic surgeries. Current guidelines rely primarily on aortic diameter for intervention; however, many dissections occur without severe dilation. Mechanical properties, such as elastic modulus have emerged as potential predictors of disease progression; nonetheless, noninvasive clinical applications remain limited. This study evaluated the relationship between the computed tomography (CT)-derived elastic modulus of the ascending aorta and the histopathological characteristics of the aortic media.

**Methods:** Thirty patients who underwent surgical ascending aorta replacement were included in this study. Preoperative CT was used to calculate the aortic elastic modulus based on geometric measurements and pulse pressure. Resected aortic specimens were subjected to histological and immunohistochemical analyses to assess elastin, collagen, vascular smooth muscle cells (VSMCs), and smoothelin expression. Correlation analyses between the CT-derived elastic modulus and aortic media composition were conducted after adjusting for age and aortic diameter.

**Results:** The CT-derived elastic modulus exhibited a significant negative correlation with elastin area and a positive correlation with collagen area. Additionally, a moderate negative correlation was observed between the elastic modulus and elastin fiber waviness. A strong negative correlation was detected between the elastic modulus and the proportion of contractile-type (smoothelin-positive) VSMCs. These findings remained significant after adjusting for confounders.

**Conclusions:** CT-derived elastic modulus of the ascending aorta reflects the underlying pathological changes, including extracellular matrix remodeling and VSMC phenotypic modulation. Noninvasive assessment of aortic mechanical properties may provide novel insights into aortic disease progression and therapeutic responses.

## Introduction

Rupture or dissection of the ascending aorta is associated with various life-threatening complications, and surgical management following onset remains highly challenging. Prophylactic surgical intervention is recommended when the risk of such events is considered high.

Current guidelines determine the threshold for surgical intervention in aortic disease primarily based on the aortic diameter, although specific criteria vary for specific aortic conditions such as bicuspid aortopathy and Marfan syndrome ^1,2^. However, the ability to predict aortic aneurysm expansion and the onset of aortic dissection remains unestablished. Studies have shown that aortic dissection frequently occurs even in patients with normal or moderately dilated aortic diameters ^3^; and no individualized treatment indicators based on patient-specific differences or aortic characteristics have been established.

In recent years, increasing research has focused on aortic mechanical properties, particularly elastic modulus, as reliable predictors of aortic disease progression, rupture risk, and dissection onset ^4-7^. Studies have demonstrated that the mechanical properties vary according to age, diameter, and aortic location.

However, despite these findings, mechanical properties have not yet been incorporated as clinical treatment indicators. One major limitation is that assessments of aortic mechanics are primarily based on resected specimens, with no effective noninvasive method available for evaluating these properties *in vivo*. To address this limitation, we previously reported the use of computed tomography (CT) as a noninvasive modality for assessing aortic mechanical properties and validated its accuracy ^8^. Moreover, changes in CT-derived mechanical properties linked with histopathological alterations of the aortic wall, such as medial degeneration and structural modifications of the extracellular matrix, have not yet been reported. Building on this foundation, the present study evaluated the relationship between the CT-derived elastic modulus and aortic histopathological characteristics to further explore its potential as a predictive tool for aortic disease progression.

## Methods

### Ethical statement

This study was approved by the Ethics Committee of Osaka University Hospital (IRB: 22026) as a non-interventional study using surplus patient specimens, and informed consent was provided by opt-out.

### Study population

We enrolled 87 consecutive surgical patients who underwent prosthetic graft replacements of thoracic aortic aneurysms at our institution between November 2020 and July 2022 (Figure 1). Ascending aortas without whole circumference or complete wall layers, patients without preoperative enhanced gated CT, and those with acute aortic dissections of the ascending aorta were excluded. Thus, 30 patients were eligible for inclusion in this study.

**Figure 1.**
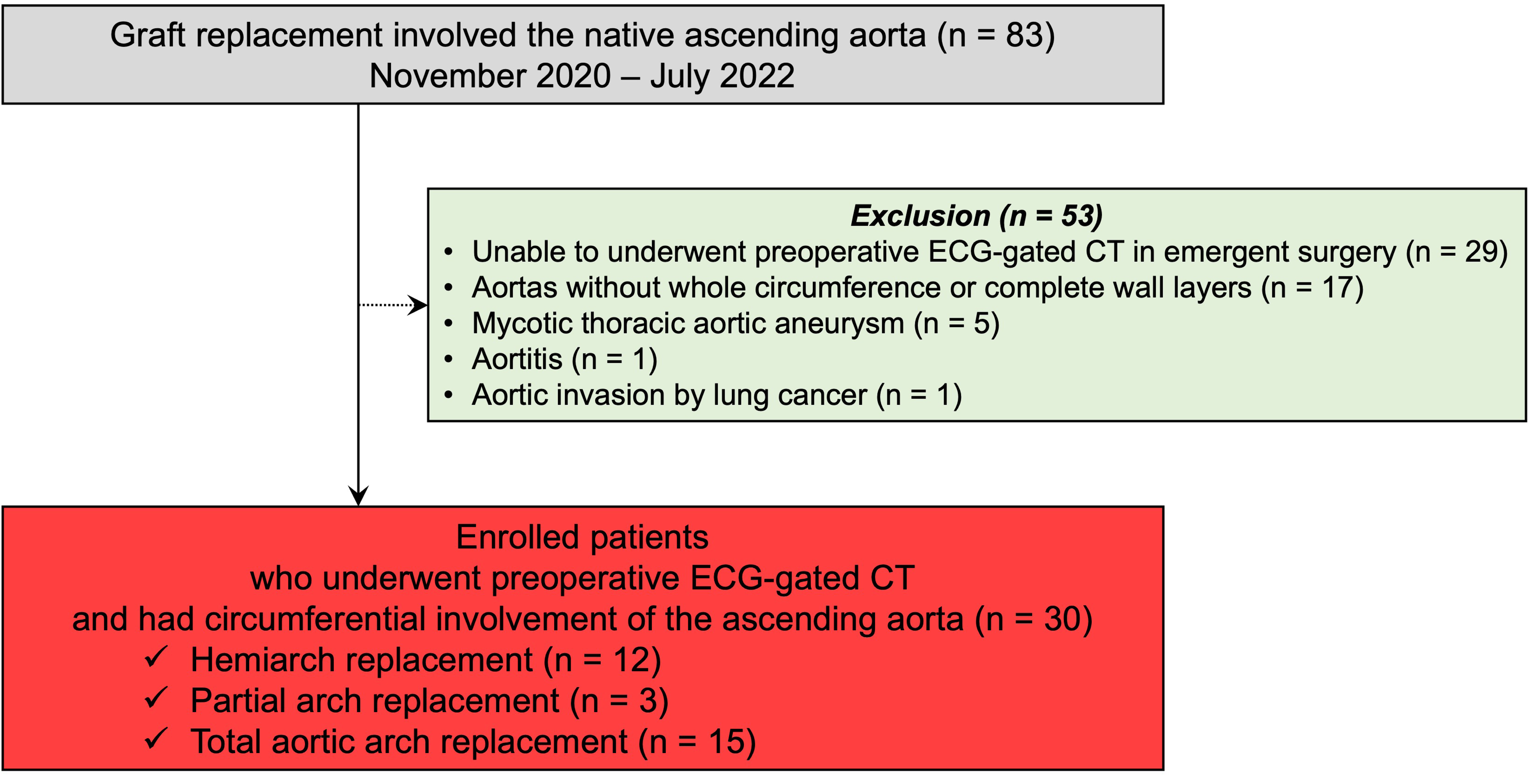
Flowchart of study population. ECG: electrocardiogram, CT: computed tomography.

### Sample preparation

All tissues of the resected ascending aorta were collected from the operating room and immediately stored at 4℃ in Ringer’s lactate solution. For the ascending aorta, specimens were received in the form of a complete ring, with the orientation marked by the surgeon. The resected ascending aorta was cut into short segments in the transverse direction, and the tissue from the anterior aspect of the greater curvature was used for histological evaluation (Supplemental Figure 1).

### Histological evaluation of resected aortic samples

Resected ascending aortic samples were fixed in 10% formalin, embedded in paraffin, and cut into 2-μm slices using a microtome for histological studies. The sections were stained with hematoxylin–eosin (HE), Picro-Sirius Red, and Elastica van Gieson (EVG).

The sections were labeled immunohistologically with polyclonal anti-SMA and polyclonal anti-smoothelin antibodies and visualized using the LSABTM kit (DAKO, K0690, Glostrup, Denmark), which is an automated immunostaining system based on the LSAB Lepto streptavidin–biotin-peroxidase method.

The images were examined by optical microscopy (KEYENCE, Osaka, Japan) and quantitative morphological analysis of each sample was performed using the Metamorph software (Molecular Devices, Sunnyvale, California, US).

The evaluation sites were determined by randomly selecting six regions, spanning from the adventitial to intimal sides that were suitable for microscopic assessment. The pathological analysis focused on the components of the aortic media, specifically the extracellular matrix components, elastin and collagen, and vascular smooth muscle cells. For elastin, an automated measurement macro developed in ImageJ software was used to determine the waviness ratio, which was calculated by automatically extracting the five longest elastin fibers from each of the six selected regions and computing the ratio of the straight-line distance between the ends of the fiber to the actual fiber length (Supplemental Figure 2). For vascular smooth muscle cells, the cellular content was assessed using αSMA staining, and their degree of differentiation was evaluated through immunohistochemistry with an anti-smoothelin antibody.

### Histopathological classification of resected aortic specimens

HE, EVG, and Sirius Red staining of the resected ascending aortic tissue were used for qualitative histopathological evaluation.

The following parameters were assessed: atherosclerosis, elastic fiber fragmentation and/or loss, elastic fiber thinning, elastic fiber disorganization, smooth muscle cell nuclear loss, smooth muscle cell disorganization, laminar medial collapse, overall medial degeneration, and medial fibrosis. The evaluation method was based on previously published criteria, and each parameter was graded on a four-point scale: none, mild, moderate, or severe ^9,10^. All specimens were evaluated by an independent, experienced histopathologist who was blinded to the clinical data.

### Calculation of CT-derived elastic modulus

We performed a preoperative electrocardiogram gated CT of the ascending aorta. The elastic moduli were calculated using the theoretical formula reported by Shirakawa et al ^8^. The formula was implemented in an analysis program connected to a medical DICOM viewer, OsiriX MD (Pixmeo SARL, Bernex, Switzerland). We used a Poisson’s ratio of 0.49, which was employed in previous studies.

We selected a cross-section of the ascending aorta, almost in the middle, between the sinotubular junction (the proximal end of the ascending aorta) and brachiocephalic artery (the distal end). Apparent calcification or atheromatous thickening was avoided. The aortic diameter was automatically calculated from the cross-sectional area extracted by region-growing segmentation of the CT images. When the aortic axis was tilted > 20°from the image plane, the images were reconstructed to the orthogonal cross-section by multiplanar reconstruction. The wall thickness was manually measured by a certified cardiovascular surgeon. The inner border of the aortic wall was the edge of the contrast-enhanced aortic lumen, and the outer border was determined based on low CT values of extra-vessel structures such as fat and the thoracic cavity. Measurements were performed at five points in the cross-section for each patient. In this case, we used the mean value as the aortic wall thickness. We determined the pulse pressure from the noninvasive blood pressure measured after the patient was placed on a scanner table before CT scanning. The von Mises stress was calculated from three-dimensional tensile stresses in the wall, assuming a symmetric and isotropic aorta.

### Statistical analysis

Continuous variables are presented as median (interquartile range [IQR]) for normal distribution, while categorical variables are presented as frequencies and percentages. The correlation between CT-derived elastic modulus values and quantitatively evaluated values of aortic media components was examined using simple linear correlation and partial correlation analyses. Correlations between aortic media components and CT-derived elastic moduli were investigated by calculating Pearson correlation coefficients and Pearson partial correlation coefficients, with adjustments for age and ascending aortic diameter as potential confounders. The cohort was divided into two groups based on median age and aortic diameter. For each group, the correlation coefficients between the aortic media components and elastic modulus were calculated after adjusting for age and aortic diameter. All p-values were two-sided, and statistical significance was set at *p*<0.050. All statistical analyses were performed using the JMP version 15.0.0 statistical software for MacOSX (SAS Institute Inc., Cary, NC, USA).

## Results

### Patient Characteristics

The baseline patient characteristics are summarized in Table 1. The median (range) patient age was 76 (49–85) years, and 75.9% (n = 22) were men. Degenerative aneurysm pathology was observed in 83.3% (n = 25) of patients, and chronic aortic dissection was observed in 16.7% (n = 5). Regarding the details of aortic pathology and surgical procedures, aortic arch aneurysm was the predominant condition (50%), with total arch replacement performed in 15 cases. Ascending aortic replacement for ascending aortic dilatation or aneurysm was also performed in 15 cases, among which 7 cases involved concomitant aortic root enlargement. The median (range) diameter of the ascending aorta on preoperative CT was 42 (35–70) mm, and the median thickness of the ascending aortic wall was 1.7 mm.

**Table 1.** Baseline characteristics of patients.

| Variables | N=30 |
| --- | --- |
| Age, yrs median (IQR) | 76 (62–78) |
| Male, n (%) | 22 (75.9) |
| BSA, m<sup>2</sup> median (IQR) | 1.62 (1.50–1.74) |
| Pathologies |  |
| Degenerative, n (%) | 25 (83.3) |
| Dissection, n (%) | 5 (16.7) |
| Medical history |  |
| Hypertension, n (%) | 20 (66.7) |
| Hyperlipidemia, n (%) | 14 (46.7) |
| Diabetes mellitus, n (%) | 6 (20.0) |
| Smoker, n (%) | 22 (73.3) |
| COPD, n (%) | 11 (36.7) |
| Coronary artery disease, n (%) | 4 (13.3) |
| Peripheral artery disease, n (%) | 4 (13.3) |
| Cancer, n (%) | 2 (6.7) |
| LVEF, % median (IQR) | 62 (53–70) |
| eGFR, mL/min/1.73m<sup>2</sup> median (IQR) | 57 (36–76) |
| FDP, µg/mL median (IQR) | 1.38 (0.34–3.82) |
| <u>Preoperative oral medication</u> |  |
| Beta blocker | 14 (46.7) |
| Statin | 15 (50.0) |
| Steroid | 0 (0.0) |
| <u>Pre-operative CT measurements</u> |  |
| Diameter of ascending aorta, mm, median (IQR) | 42 (36–49) |
| Thickness of aortic wall, mm, median (IQR) | 1.7 (1.5–1.8) |
| Maximum aneurysm diameter, mm, median (IQR) | 55 (53-63) |
| Aortic valve characteristics |  |
| Tricuspid aortic valve | 25 (30.0) |
| Bicuspid aortic valve | 5 (16.7) |
| Severe aortic valve stenosis | 2 (6.7) |
| Severe aortic valve regurgitation | 7 (23.3) |

BSA, body surface area; COPD, chronic obstructive pulmonary disease; CT, computed tomography; LVEF, left ventricular ejection fraction; eGFR, estimated glomerular filtration rate; IQR, interquartile range.

### Histopathological classification of the ascending aorta specimens

The histopathological characteristics of the ascending aorta samples are summarized in Table 2. Of the 30 analyzed aortic specimens, 46.7% showed no atherosclerosis, while the remaining samples demonstrated varying severity (mild: 6.7%; moderate: 26.6%; severe: 20.0%). Elastic fiber fragmentation and/or loss was present in the majority of cases, with 63.3% graded as mild. Elastic fiber thinning was also frequently observed (63.4%), with 40.0% graded as mild. Disorganization of elastic fibers was rare (10.0%). Loss of smooth muscle cell nuclei was noted in 73.3% of cases, whereas disorganization was not observed. Laminar medial collapse was absent in 73.3% and, when present, was limited to mild or moderate changes. Overall medial degeneration was classified as mild or moderate in 83.3% and 6.7% of cases, respectively. Medial fibrosis was universally observed, with moderate or severe involvement in 63.3% of cases.

**Table 2.** Pathological characteristics of ascending aorta samples.

|  |  |
| --- | --- |
|  | N = 30 |
| <u>*Atherosclerosis, n (%)*</u> |  |
| none | 14 (46.7) |
| mild | 2 (6.7) |
| moderate | 8 (26.6) |
| severe | 6 (20.0) |
| <u>*Elastic fiber fragmentation and/or loss, n (%)*</u> |  |
| none | 1 (3.3) |
| mild | 19 (63.3) |
| moderate | 6 (20.0) |
| severe | 1 (3.3) |
| unevaluable | 3 (10.0) |
| <u>*Elastic fiber thinning, n (%)*</u> |  |
| none | 11 (36.6) |
| mild | 12 (40.0) |
| moderate | 2 (6.7) |
| severe | 2 (6.7) |
| unevaluable | 3 (10.0) |
| <u>*Elastic fiber disorganization, n (%)*</u> |  |
| none | 24 (80.0) |
| mild | 1 (3.3) |
| moderate | 2 (6.7) |
| severe | 0 (0.0) |
| unevaluable | 3 (10.0) |
| <u>Smooth muscle cell nuclei loss, n (%)</u> |  |
| none | 0 (0.0) |
| mild | 22 (73.3) |
| moderate | 3 (10.0) |
| severe | 0 (0.0) |
| unevaluable | 3 (10.0) |
| <u>Smooth muscle cell disorganization, n (%)</u> |  |
| none | 27 (90.0) |
| mild | 0 (0.0) |
| moderate | 0 (0.0) |
| severe | 0 (0.0) |
| unevaluable | 3 (10.0) |
| <u>Laminar medial collapse, n (%)</u> |  |
| none | 22 (73.3) |
| mild | 2 (6.7) |
| moderate | 3 (10.0) |
| severe | 0 (0.0) |
| unevaluable | 3 (10.0) |
| <u>Overall medial degeneration, n (%)</u> |  |
| none | 0 (0.0) |
| mild | 25 (83.3) |
| moderate | 2 (6.7) |
| severe | 0 (0.0) |
| unevaluable | 3 (10.0) |
| <u>Medial fibrosis, n (%)</u> |  |
| none | 0 (0.0) |
| mild | 8 (26.7) |
| moderate | 11 (36.6) |
| severe | 8 (26.7) |
| unevaluable | 3 (10.0) |

### Correlation between aortic media components and CT-derived elastic modulus of the ascending aorta by computed tomography

The correlation between the quantitative values of the ascending aortic media components and CT-derived elastic modulus is shown in Figures 2 and 3. A negative correlation was observed between the elastin area and elastic modulus (*r*² = 0.75, *p* < .01), whereas the collagen area exhibited a positive correlation with elastic modulus (*r*² = 0.61, *p* < .01). No significant correlation was found between the vascular smooth muscle cell area and elastic modulus (*r*² = 0.02). Morphological evaluation of the media revealed that the degree of waviness of the elastin fibers decreased as the elastic modulus increased (*r*² = 0.27, *p* < .01). Regarding the differentiation of vascular smooth muscle cells, both the area of contractile-type smooth muscle cells and their proportion relative to the total smooth muscle cell population decreased with increasing elastic modulus (*r*² = 0.74, *p* < .01).

**Figure 2.**
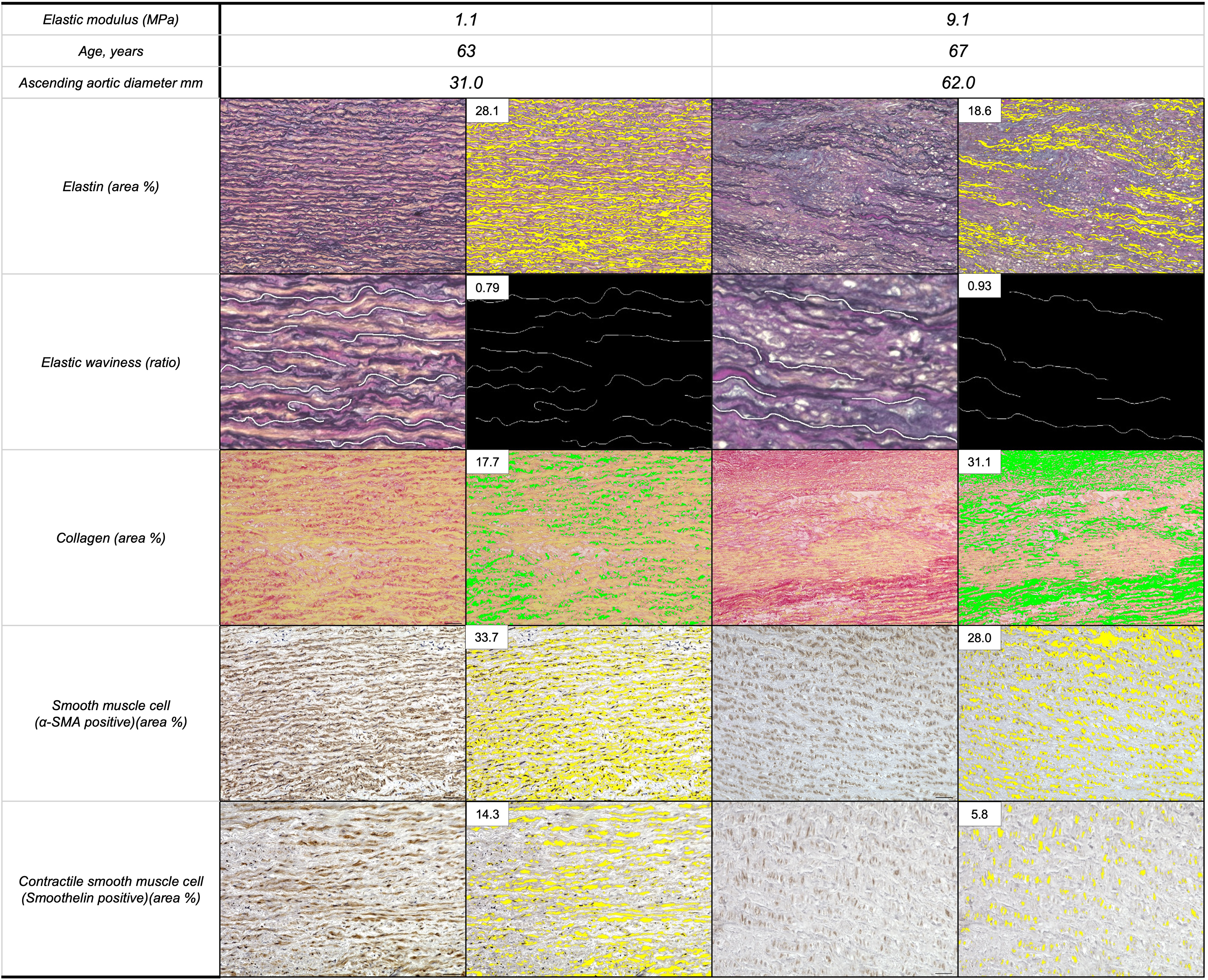
Representative examples of changes in CT-derived ascending aortic elastic modulus and corresponding histological evaluation images. Scale bar: 50 μm. CT: computed tomography; SMA: smooth muscle actin

**Figure 3.**
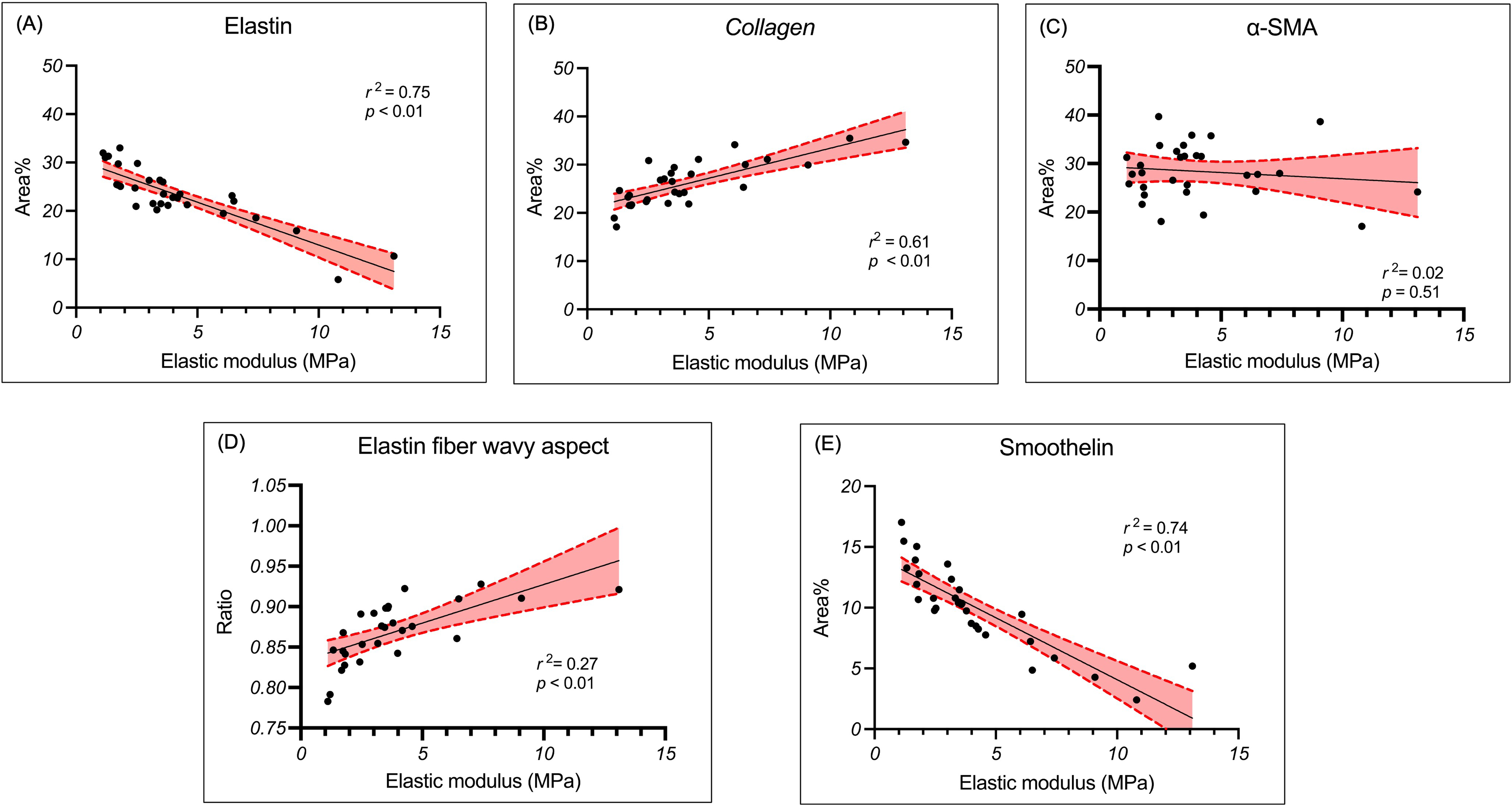
Correlation of aortic media components with elastic modulus of ascending aorta in computed tomography calculation. SMA: smooth muscle actin

### Correlation between aortic media composition, elastic modulus, and cofounding variables

The correlation between aortic media composition, elastic modulus, and confounding variables was determined using a correlation analysis adjusted for age and ascending aortic diameter (Table 3). The elastic modulus was significantly correlated with the areas of elastin (*r*² = 0.78, *p* < .01), collagen (*r*² = 0.72, *p* < .01), smooth muscle cells (*r*² = 0.43, p < .01), smoothelin-positive cells (*r*² = 0.82, *p* < .01), and the waviness ratio of elastin fibers (*r*² = 0.68, *p* < .01) after adjusting for age and ascending aortic diameter. Additionally, in each category of age and ascending aortic diameter, the elastin, collagen, and smoothelin-positive areas, and waviness ratio of elastin fibers exhibited a significant correlation with the CT-derived elastic modulus, even after adjusting for age and ascending aortic diameter.

**Table 3.** Regression coefficients of aortic medial components factor for ascending aortic elastic modulus measured by computer tomography.

| Variables | Univariable |  | Age and ascending aortic diameter-adjusted |  |
| --- | --- | --- | --- | --- |
| | $r^2$ | $p$ - values | $r^2$ | $p$ - values |
| Total cohort |  |  |  |  |
| Elastin | 0.75 | < .01 | 0.78 | < .01 |
| Collagen | 0.61 | < .01 | 0.72 | < .01 |
| Smooth muscle cell | 0.02 | 0.51 | 0.43 | < .01 |
| Smoothelin (Contractile type) | 0.74 | < .01 | 0.82 | < .01 |
| Waviness ratio of elastin fibers | 0.27 | < .01 | 0.68 | < .01 |
| Old-age category (76–85 years) |  |  |  |  |
| Elastin | 0.76 | < .001 | 0.76 | < .01 |
| Collagen | 0.45 | < .01 | 0.65 | < .01 |
| Smooth muscle cell | 0.22 | 0.08 | 0.39 | 0.13 |
| Smoothelin (Contractile type) | 0.63 | < .01 | 0.87 | < .01 |
| Waviness ratio of elastin fibers | 0.14 | 0.19 | 0.66 | 0.01 |
| Young-age category (49–75 years) |  |  |  |  |
| Elastin | 0.80 | < .01 | 0.84 | < .01 |
| Collagen | 0.75 | < .01 | 0.82 | < .01 |
| Smooth muscle cell | 0.03 | 0.57 | 0.6 | 0.05 |
| Smoothelin (Contractile type) | 0.82 | < .01 | 0.83 | < .01 |
| Waviness ratio of elastin fibers | 0.46 | 0.01 | 0.78 | < .01 |
| Large-diameter of ascending aorta |  |  |  |  |
| category (43–70 mm) |  |  |  |  |
| Elastin | 0.81 | < .01 | 0.94 | < .01 |
| Collagen | 0.75 | < .01 | 0.93 | < .01 |
| Smooth muscle cell | 0.04 | 0.46 | 0.79 | < .01 |
| Smoothelin (Contractile type) | 0.77 | < .01 | 0.94 | < .01 |
| Waviness ratio of elastin fibers | 0.14 | 0.19 | 0.91 | < .01 |
| Small-diameter of ascending aorta |  |  |  |  |
| category (35–42 mm) |  |  |  |  |
| Elastin | 0.61 | < .001 | 0.70 | < .01 |
| Collagen | 0.52 | < .01 | 0.57 | 0.01 |
| Smooth muscle cell | 0.01 | 0.69 | 0.29 | 0.26 |
| Smoothelin (Contractile type) | 0.79 | < .01 | 0.82 | < .01 |
| Waviness ratio of elastin fibers | 0.69 | < .01 | 0.73 | < .01 |

## Discussion

The results of this study demonstrate a correlation between the noninvasive CT-derived elastic modulus and extracellular matrix composition, as well as vascular smooth muscle cell content within the ascending aortic media. Furthermore, morphological changes in elastin and the vascular smooth muscle cell phenotype, specifically the waviness ratio of elastin fibers and degree of vascular smooth muscle cell differentiation (contractile type), also showed a significant correlation.

Previous studies have evaluated the main extracellular matrix components of the aortic media, elastin and collagen, which largely determine the aortic elastic modulus. In all reported results, an increase in the elastic modulus of resected aortic specimens was negatively correlated with elastin content and positively correlated with collagen content ^7,11^. Furthermore, the influence of age and aortic diameter on tissue composition has also been reported ^8,11^. Here, the CT-derived elastic modulus exhibited a similar correlation with both extracellular matrix and vascular smooth muscle cell content. Regarding the morphological evaluation of elastin fibers, previous studies measured the waviness of the elastic lamellae in the aorta of diabetic mouse models using Synchrotron X-ray micro-CT exploration. These studies suggest that measurement of elastin waviness may serve as an indicator of aortic elasticity ^12^.

While previous studies have reported no significant differences in the proportion of vascular smooth muscle cells between normal and aneurysmal aortas ^7^, our findings were consistent in showing no correlation between the CT-derived elastic modulus and overall vascular smooth muscle cell content. However, an important and novel observation in our study was the significant correlation between the CT-derived elastic modulus and both the absolute area and proportion of smoothelin-positive smooth muscle cells, which are indicative of a contractile phenotype. This suggests that the mechanical properties of the aorta may be more closely associated with the functional state of smooth muscle cells rather than total quantity.

Previous studies by Honma et al. suggested that the regulation of smoothelin may be involved in the development and progression of aortic disease based on pathological image analysis comparing normal human aorta and aortic aneurysm samples. They reported a marked decrease in smoothelin expression in α-smooth muscle actin-positive vascular smooth muscle cells of the aortic media, making it difficult to identify positive cells by immunohistochemistry ^13^. In addition, differences in smoothelin expression in the ascending aorta have been reported based on aortic valve morphology ^14^. These previous reports suggest that smoothelin regulation may be involved in the development and progression of aortic diseases. It has also been reported that the expression of smooth muscle contractile marker genes, such as *MYH11*, *TAGLN*, *CNN1*, and *SMTN*, is downregulated on stiff substrates. Conversely, genes associated with the proliferation and synthetic phenotype of smooth muscle cells are upregulated on stiff substrates compared with those on soft substrates ^15,16^. A common finding between our results and those of previous studies is that increased tissue stiffness, reflected by a higher elastic modulus, was associated with reduced expression of contractile-type smooth muscle cell genes. Therefore, our findings capture a similar trend to that reported in previous studies.

The clinical significance of this study lies in the suggestion that noninvasive CT-derived measurements of the ascending aortic elastic modulus may serve as useful parameters for capturing changes in the tissue environment of the ascending aorta. The noninvasive CT-derived aortic elastic modulus holds potential as a novel tissue-based indicator for assessing the progression of aortic aneurysms or the onset of aortic dissection. Furthermore, it may offer a means to noninvasively and accurately evaluate tissue changes and the effectiveness of pharmacological therapies targeting aortic pathology. Further research is needed to clarify the relationship between CT-derived aortic mechanical properties and the tissue environment, as well as their long-term changes and associations with the development and progression of aortic diseases.

This study has several limitations. First, the sample size, including the number of resected aortic specimens, was relatively small. As a result, the findings may be subject to selection bias, potentially limiting their generalizability to broader patient populations. Second, most of the specimens were derived from patients with atherosclerotic aortic disease. While this enhances the relevance of our findings to degenerative aneurysm mechanisms, it remains uncertain whether the results can be extrapolated to patients with genetically triggered aortic disorders, such as Marfan syndrome, which were not represented in this cohort. Similarly, the applicability of these findings to valvular aortopathy is limited due to the small number of such cases included. Further studies are warranted to validate these observations in genetically mediated and non-atherosclerotic aortic diseases. Third, histological analysis was limited to the central anterior region of the ascending aorta. While the CT-based mechanical model assumes a homogeneous circular geometry, the ascending aorta is anatomically heterogeneous and often exhibits localized atherosclerotic remodeling. Therefore, regional variations across the full aortic circumference may not have been fully captured, potentially underestimating the extent of structural diversity.

## Conclusion

this study identified significant associations between the CT-derived elastic modulus of the ascending aorta and underlying histopathological features, including elastic fiber waviness and vascular smooth muscle cell phenotype. These findings suggest that CT-based mechanical assessment may serve as a noninvasive surrogate marker of medial degeneration and could contribute to improved risk stratification for ascending aortic enlargement and dissection. Further studies are warranted to validate these results in broader patient populations and to clarify their potential clinical utility.

## Acknowledgements

We thank Editage for editing a draft of this manuscript.

## Materials and Data Availability

The data underlying this article will be shared on reasonable request to the corresponding author.

## Figure Legends

**Supplemental Figure 1.** Workflow for ascending aortic tissue preparation for histological analysis.

(A) Appearance of the resected ascending aortic specimen showing the greater curvature orientation. The proximal and distal portions of the ascending aorta were identified and marked; the yellow arrow indicates the transection site.

(B) The specimen was opened longitudinally along the lesser curvature (yellow dotted arrow) and laid flat. The region of the greater curvature designated for histological analysis is outlined with black dotted lines. The red arrow indicates the direction of antegrade blood flow.

(C) The marked region was sectioned into parallel longitudinal strips (black dotted box) for paraffin embedding and histological processing.

(D) Representative haematoxylin and eosin-stained cross-section. The central analysis area was defined as a 3 × 3 grid (dotted box) for quantitative morphological evaluation. Scale bar: 200 μm.

**Supplemental Figure 2.** Quantification method of elastin fiber waviness in the aortic media.

(A) Representative elastin-stained histological section of the aortic media using Elastica van Gieson staining, demonstrating the architecture of the medial elastic lamellae.

(B) High-magnification image showing individually segmented elastic fibers, sequentially numbered for waviness analysis.

(C) Binary image illustrating the morphological isolation of each elastic fiber following segmentation.

(D) Schematic representation of waviness ratio calculation. The yellow line indicates the straight-line distance between the endpoints of a fiber, and the white line indicates the actual curved fiber path. Waviness ratio is defined as the straight length divided by the curved length. the straight length divided by the wave (curved) length.

